# Resting calcium ion fluxes protect cells from fast mitochondrial fragmentation, cell stress responses, and immediate transcriptional reprogramming

**DOI:** 10.1101/2025.02.06.636834

**Authors:** Caroline Fecher, Annemarie Sodmann, Felicitas Schlott, Juliane Jaepel, Franziska Schmitt, Isabella Lengfelder, Thorsten Bischler, Bernhard Nieswandt, Konstanze F. Winklhofer, Robert Blum

## Abstract

Homeostatic calcium ion (Ca^2+^) fluxes between the endoplasmic reticulum, cytosol, and extracellular space occur not only in response to cell stimulation but also in unstimulated cells. Using murine astrocytes as a model, we asked whether there is a signaling function of these resting Ca^2+^-fluxes. The data showed that endoplasmic reticulum (ER) Ca²⁺ depletion, induced by sarcoplasmic/endoplasmic reticulum Ca²⁺-ATPase (SERCA) inhibition, resulted to prolonged Ca²⁺ influx and mitochondrial fragmentation within 10 to 30 minutes. This mitochondrial fragmentation could be prevented in Ca^2+^- free medium or by inhibiting store-operated Ca^2+^ entry (SOCE). Similarly, attenuation of STIM proteins, which are vital ER Ca^2+^ sensors, protected mitochondrial morphology. On the molecular level, ER Ca^2+^ depletion, achieved either by removing extracellular Ca^2+^ or through acute SERCA inhibition, led to changes in gene expression of about 13% and 41% of the transcriptome within an hour, respectively. Transcriptome changes were associated with universal biological processes such as transcription, differentiation, or cell stress. Strong increase in expression was observed for the transcription factor ATF4, which is under control of the kinase PERK (EIF2AK3), a key protein involved in ER stress. Corroborating these findings, PERK was rapidly phosphorylated in Ca^2+^-free medium or after acute pharmacological inhibition of SOCE. In summary, resting, homeostatic Ca^2+^ fluxes prevent immediate-early cell stress and transcriptional reprogramming.

## Introduction

All eukaryotic cells balance their calcium ion (Ca^2+^) fluxes to maintain cell homeostasis (*1, 2*). Recent research in neural cells demonstrated that baseline cellular Ca^2+^ levels are managed by homeostatic Ca^2+^ fluxes between the ER Ca^2+^ store and the extracellular space (*3–7*). These homeostatic, resting Ca^2+^ fluxes are initially triggered by an evolutionary conserved process by which passive Ca^2+^ leakage from the ER store (*5, 8*) results in a surprisingly strong intracellular Ca^2+^ flux (*4, 5, 8–10*). To counterbalance the inevitable passive loss of Ca^2+^ from the ER, Ca^2+^ is actively pumped back into the ER lumen by the sarcoplasmic/endoplasmic reticulum Ca^2+^-ATPase (SERCA). In neurons, much of the Ca^2+^ lost through ER leakage is actively transported to the extracellular space and replaced via homeostatic Ca^2+^ influx mechanisms (*3, 6, 7, 11*). Therefore, inhibitors of store operated Ca^2+^ entry (SOCE) can cause an immediate neuronal ER Ca^2+^ underload (*3, 12*). This effect can be measured in about 25% of cultured astrocytes within minutes after SOCE inhibition (*4, 5*). Hence, neural cells employ energy intensive Ca^2+^ fluxes to balance resting Ca^2+^ levels and to avoid ER Ca^2+^ underload for maintaining cell signaling and health (*2, 13*).

The concept of Ca^2+^ overload, which refers to increased Ca^2+^ levels in the cytosol, mitochondria, and/or the ER stores, has garnered significant attention. In this concept, excitotoxic Ca^2+^ influx, which can occur after stroke or during epileptic seizures, causes oxidative stress, mitochondrial dysfunction, and ER stress (*14*). Ca^2+^ overload may also be important in slowly progressing neurodegenerative diseases like in Alzheimer’s (*15*) or Parkinson’s disease (*16*). However, the role of ER Ca^2+^ depletion to cell health and pathology is less studied, even though it induces long-lasting Ca^2+^ influx from the extracellular space (*4*). ER Ca^2+^ depletion is rapidly sensed by stromal interaction molecule proteins (STIM1 and STIM2) and ER stress sensors such as PERK (PKR-Like ER Kinase), a key protein in ER quality control pathways (*17*). These ER stress responses are pivotal in regulating both pro-apoptotic and anti-apoptotic pathways, balancing between cell death and cell survival (*17, 18*).

In this study, we used cultured astrocytes as a highly responsive cell model for Ca^2+^ signaling, to investigate signaling functions of homeostatic Ca^2+^ fluxes.

## Results

### ER Ca^2+^ depletion causes prolonged SOCE with a transient increase in mitochondrial Ca^2+^

We recently described the Ca^2+^ signaling repertoire and toolkit of murine cortical astrocytes in culture (*4*). In these cells, the ER Ca^2+^ leak is counteracted by the SERCA protein and homeostatic SOCE (outlined in Fig. 1A). To investigate the ER Ca^2+^ leak in these cells, we examined the effects of blocking SERCA. Fast perfusion with the SERCA blocker thapsigargin induced rapid ER Ca^2+^ depletion and sustained SOCE (Fig. 1B, data were re-plotted for didactic reasons. Original data from (*4*)).

**Figure 1.**
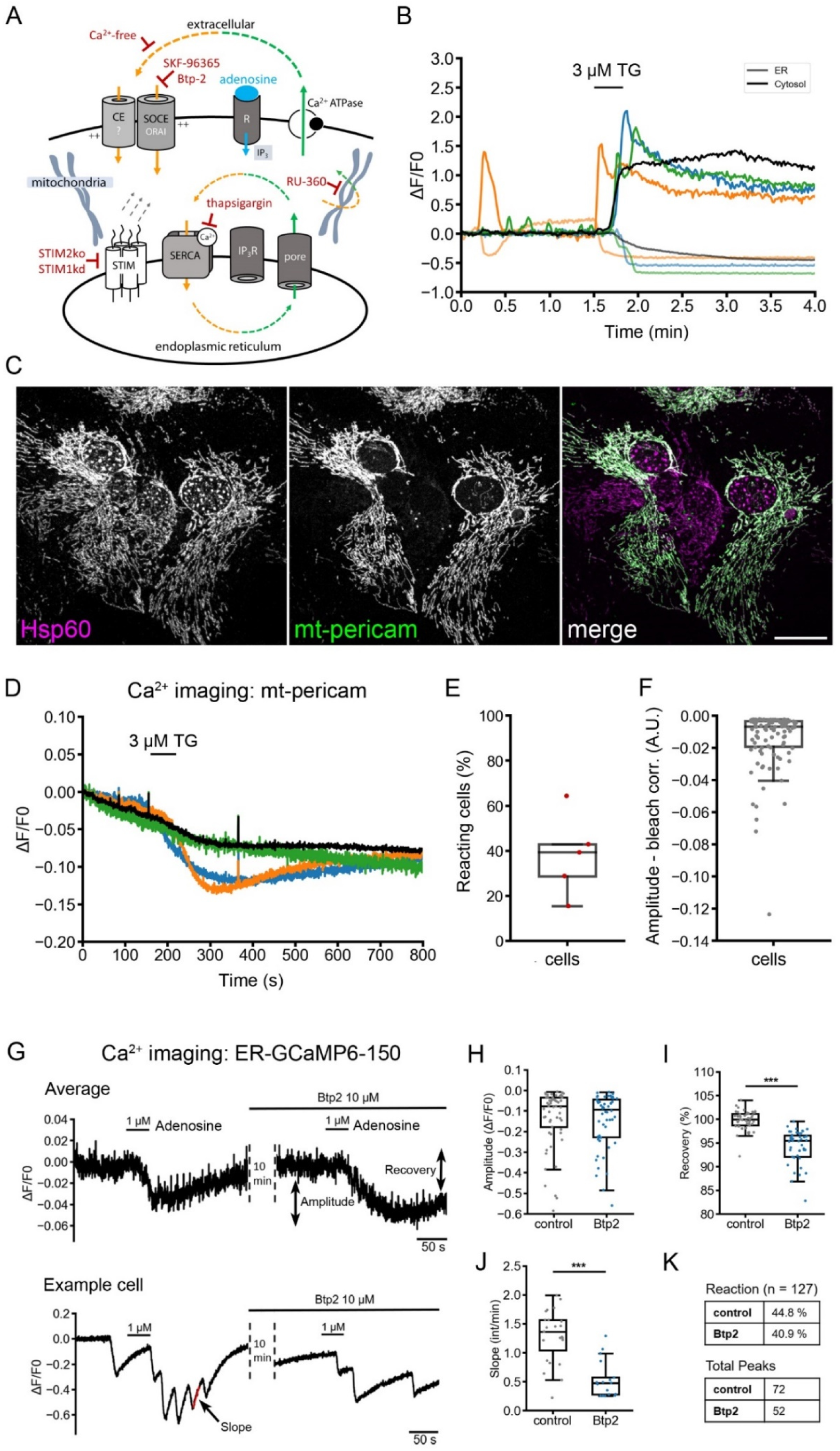
Ca^2+^ dynamics in response to SERCA blockage in astrocytes. **A.** Outline of resting Ca^2+^ fluxes in astrocytes. At rest, Ca^2+^ is also lost to the extracellular space, which is compensated by resting Ca^2+^ entry. **B.** Immediate Ca^2+^ responses to SERCA blockage. 3 µM thapsigargin was added for 20 s while ER (ER- GCaMP6-150) and cytosolic Ca^2+^ (Cal590) were imaged. Data is plotted as relative change in fluorescence (deltaF/F_0_) over time. In grey: average signal for the ER Ca^2+^ signal; in black: average signal for the cytosolic Ca^2+^ signal; n = 31 cells from a representative experiment. Three single cell examples are shown for both conditions (data taken from (*4*)). **C.** Lentiviral expression of mt-pericam in astrocytes. Representative confocal image of mt-pericam, labeled against GFP and the mitochondrial marker Hsp60. Note the tubular structure of the mitochondria. Scale bar: 20 µm. **D.** Acute SERCA blockage causes only transient and brief Ca^2+^ responses in mitochondria expressing mt-pericam. Data is plotted as relative change in fluorescence (deltaF/F_0_) over time. Shown are signal traces of three single cells (blue, green, orange) and an average signal trace (black) of multiple individual cells (data from 5 measurements using 3 independent cultures). **E.** Graph showing percentage of glial cells with mt-pericam responses to acute SERCA blockade. Single data points (dark red dots) indicate mean values from 5 measurements. About 40% of all cells show Ca^2+^ influx responses mediated by mt-pericam. The mean and SD are indicted in the boxplot. **F.** Graph presents signal amplitude values of mt-pericam after acute SERCA blockade. Data represent 104 single cells (grey dots) from 5 measurements (3 cultures). The mean value is indicted in the boxplot. **G.** Refilling of ER calcium is delayed in the presence of the ORAI blocker Btp2. ER calcium was monitored using ER-GCaMP6-150. Cells were stimulated with 1 µM adenosine. Then cells were incubated in Btp2 for 10 minutes and the stimulation was repeated. Data is plotted as relative change in fluorescence (deltaF/F_0_) over time and represents the average signal of n = 13 cells from one representative measurement. Signal amplitude (indicated) and ER Ca^2+^ signal recovery (indicated) were computed. As many cells oscillate in response to adenosine (*4*), the slope of the signal recovery was computed as well. **H-K. Ca^2+^ signal analysis.** Graphs show quantification of relative signal amplitude (ΔF/F_0_) (in H), signal recovery (in %, in I), slope (relative intensity per minute, in J), and percentual reactivity to 1 µM adenosine (in K). Results from single cells are indicated as dots, mean values and SD in the boxplots. Amplitude: n = 127, not normally distributed (Shapiro), Mann Whitney U-Test p = 0.35; Recovery: n = 109, not normally distributed (Shapiro), Mann Whitney U-Test p < 0.001; Slope: control: n = 65, Btp2 = 42, not normally distributed (Shapiro), Mann Whitney U-Test p < 0.001.

High levels of Ca^2+^ in the cytosol can be buffered by mitochondria. Therefore, we asked whether this prolonged SOCE leads to a corresponding increase in mitochondrial Ca^2+^ levels. To explore this, we expressed the genetic Ca^2+^ indicator mt-pericam (*19*), which is localized to the mitochondrial matrix, using lentiviral vectors. Mt-pericam specifically targeted mitochondria, which exhibited their characteristic elongated network (Fig. 1C). We imaged mt-pericam via excitation/emission at 380/520 nm and at a frequency of 5 Hz. Under these conditions, a rise in mitochondrial Ca^2+^ was indicated by a decrease in fluorescence intensity (Fig. 1D). Indeed, thapsigargin induced a transient increase in mitochondrial Ca^2+^ in about 40% of transduced cells (Fig. 1D, E). The amplitude of this change varied between individual cells (Fig. 1F). The data show that ER Ca^2+^ depletion by SERCA inhibition causes a sustained Ca^2+^ influx from the extracellular space, while mitochondrial Ca^2+^ levels are only transiently affected over a period of about five minutes.

We next examined ER Ca^2+^ dynamics in response to inhibition of stimulated SOCE. In astrocytes, adenosine functions as a metabotropic agonist of ER Ca^2+^ release through the adenosine receptor Adora1 (*4*). After adenosine stimulus, Btp2, an inhibitor of SOCE (*20, 21*), has been shown to delay the re-filling of the ER Ca^2+^ store (*4*). Based on these observations, we reasoned that acute SOCE inhibition should induce transient ER Ca^2+^ underload. To test this hypothesis, we monitored ER Ca^2+^ levels using the genetic ER Ca^2+^ sensor ER-GCamp6-150 (*22*) in the presence of extracellular Ca^2+^. Treatment of astrocytes with 1 µM adenosine caused a rapid drop in ER Ca^2+^ levels, followed by subsequent ER refilling (Fig. 1G). Next, cells were perfused with 10 µM Btp2 for 10 minutes, a condition known to induce ER Ca^2+^ underload in neurons (*3*). When cells were re-stimulated with 1 µM adenosine, they exhibited a typical ER Ca^2+^ release amplitude in responsive cells (Fig. 1G, H), but showed delayed recovery of the ER Ca^2+^ signal, suggesting delayed refilling (Fig. 1I). In oscillating cells, this effect could be seen by the reduced slope (Fig. 1J). Moreover, in oscillating cells, Btp2-preincubation reduced the percentual reactivity to adenosine stimulus and the number of total Ca^2+^ signal peaks (Fig. 1K).

Thus, the direct ER Ca^2+^ imaging reveals that the Btp-2 sensitive SOCE prevents ER Ca^2+^ underload in unstimulated cells and supports ER refilling with Ca^2+^ from extracellular after stimulation.

### Sustained SOCE induces mitochondrial fragmentation

Under SERCA blockage, long-lasting increase in cellular Ca^2+^ was observed in the cytosol; however, this was not accompanied by a similar increase in mitochondrial Ca^2+^ (Fig. 1). Previously, ER Ca^2+^ depletion triggered mitochondrial fragmentation in liver epithelial cells (*23*). Therefore, we asked whether and when mitochondrial fragmentation occurs in astrocytes after SERCA blockage, and if the subsequent long-lasting SOCE is upstream of mitochondrial fragmentation.

Astrocytes were loaded with Mitotracker CMXros, a membrane potential-dependent fluorescent dye that accumulates in mitochondria. We then imaged cells under continuous perfusion in medium containing extracellular Ca^2+^ and blocked SERCA with 3 µM thapsigargin (Fig. 2A). To quantify organelle fragmentation based on their 2D particle size (Fig. S1, Fig. 2B, C), we established a size range of 5-15 pixel^2^ to identify fragmented mitochondria (Fig. 2C; DMSO, mean ± SD = 42.9 ± 1.7; thapsigargin, mean ± SD = 130.7 ± 4.2). Mitochondrial fragmentation was visible within 10 minutes and peaked after 30 minutes (Fig. 2B).

**Figure 2.**
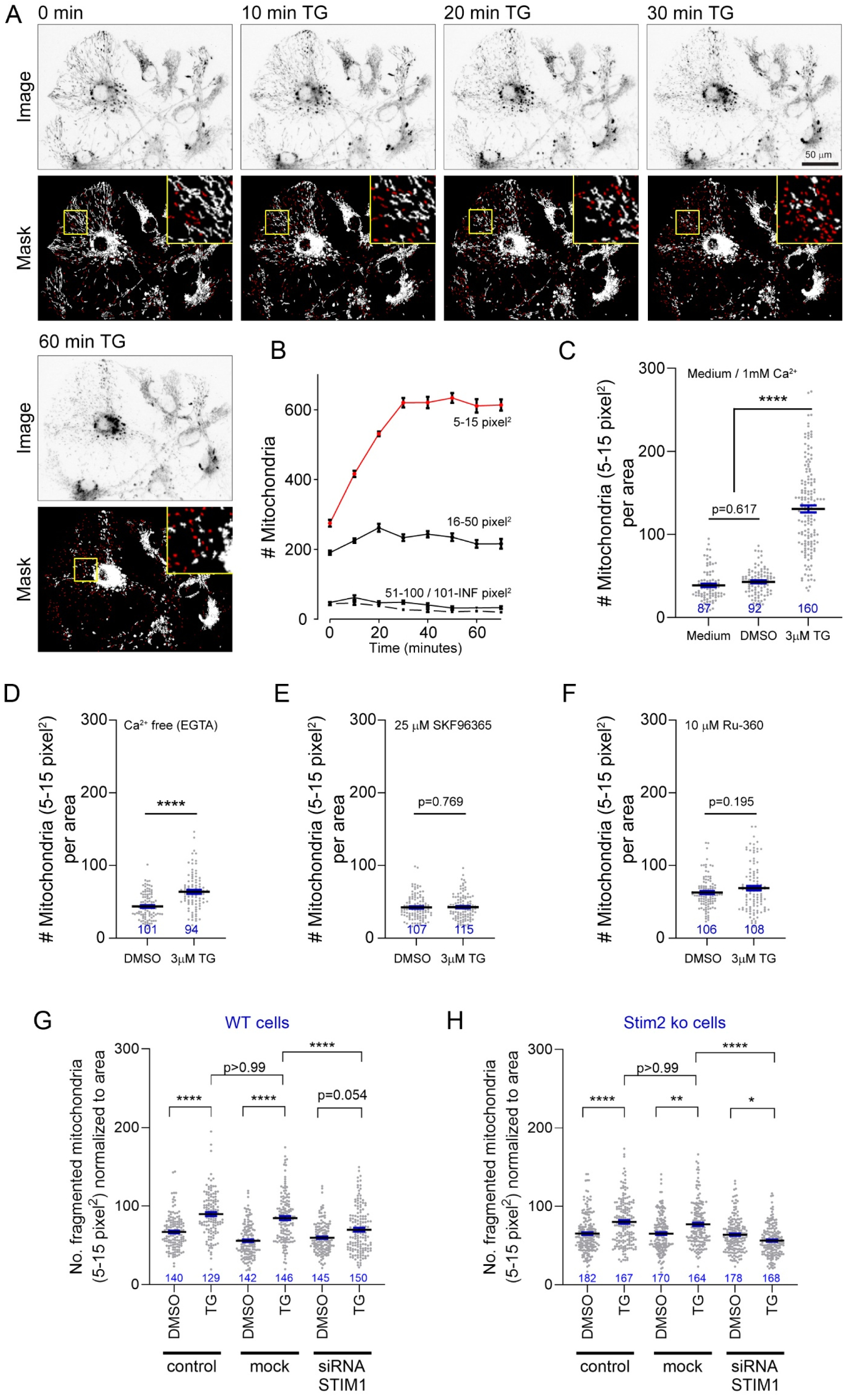
Fast mitochondrial fragmentation is induced by SOCE in response the ER Ca^2+^ depletion. Live imaging of mitochondrial morphology after permanent ER Ca^2+^ depletion with thapsigargin (TG) or the solvent control (DMSO). **A.** Time series of live cell images of mitochondria in astrocytes. Cells were labelled with the dye CMXros and imaged by fluorescence microscopy. Segmentation masks were used to quantify mitochondrial fragmentation. **B.** Mitochondrial particle quantification. CMXros-positive particles were categorized in four groups depending on their size in the image masks. Size normalization is expressed in pixel² in accordance with the raster graphics of the bioimage dataset. Mitochondrial fragmentation is best described by an increase in ‘small’ CMXros-positive particles (in red, 5 - 15 pixel²). **C.** Thapsigargin-induced fragmentation of mitochondria in astrocytes. Quantification of fragmented mitochondria (particle size 5 - 15 pixel^2^) per area after 30 minutes incubation in medium (1 mM CaCl_2_; n = 87), with vehicle (DMSO; n = 92), or thapsigargin (3 μM, TG; n = 160). Kruskal-Wallis test (H(3) = 214, P < 0.0001) with Dunn’s multiple comparison test. **D.** Thapsigargin-induced fragmentation of mitochondria in Ca^2+^ free medium. Quantification of fragmented mitochondria (particle size 5 - 15 pixel^2^) per area after 30 minutes incubation in medium supplemented with 0.1 mM EGTA and vehicle (DMSO; n = 101), or thapsigargin (3 μM, TG; n = 94). Kolmogorov-Smirnov test (D = 0.39, p < 0.0001). **E.** Thapsigargin-induced fragmentation of mitochondria in medium supplemented with 25μM SKF96365. Quantification of fragmented mitochondria (particle size 5-15 pixel^2^) per area after 30 minutes incubation in medium supplemented with 25μM SKF96365 and vehicle (DMSO; n=107), or thapsigargin (3 μM, TG; n = 115). Kolmogorov-Smirnov test (D = 0.09, p = 0.77). **F.** Thapsigargin-induced fragmentation of mitochondria in medium supplemented with 10μM Ru-360. Quantification of fragmented mitochondria (particle size 5 - 15 pixel^2^) per area after 30 minutes incubation in medium supplemented with 10 μM Ru-360 and vehicle (DMSO; n = 106), or thapsigargin (3 μM, TG; n = 108). Kolmogorov-Smirnov test (D = 0.15, p = 0.20). **G.** Thapsigargin-induced fragmentation of mitochondria in WT astrocytes under Stim1 knockdown. Astrocytes were treated with lipofectamine (ctr), mock siRNA (mock) or siRNA against Stim1 and incubated with medium plus vehicle (DMSO; n ≥ 140) or thapsigargin (3 μM, TG; n ≥ 129) for 30 minutes. Quantification of fragmented mitochondria (particle size 5 - 15 pixel^2^) per area. Kruskal-Wallis test (H(6)=160.6, p < 0.0001) with Dunn’s multiple comparison test. **H.** Thapsigargin-induced fragmentation of mitochondria in Stim2^-/-^ astrocytes under Stim1 knockdown. Astrocytes deficient in Stim2 (ko) were treated with lipofectamine (ctr), mock siRNA (mock) or siRNA against Stim1 and incubated with medium plus vehicle (DMSO; n ≥ 170) or thapsigargin (3 μM, TG; n ≥ 164) for 30 minutes. Quantification of fragmented mitochondria (particle size 5 - 15 pixel^2^) per area. Kruskal-Wallis test (H(6) = 86.39, p < 0.0001) with Dunn’s multiple comparison test.

Next, cells were treated with thapsigargin under Ca^2+^ free conditions (EGTA; Fig. 2D), which reduced the extent of mitochondrial fragmentation seen after 30 minutes (DMSO, mean ± SD = 43.6 ± 1.9; TG, mean ± SD = 63.8 ± 2.7; p<0.0001, Kolmogorov Smirnov test). Moreover, mitochondrial fragmentation was entirely blocked under SKF-96365, a SOCE inhibitor, (Fig. 2E; DMSO, mean ± SD = 42.4 ± 1.6; TG, mean ± SD = 41.2 ± 1.6; p=0.7695, Kolmogorov Smirnov test) and by RU-360, a mitochondrial Ca^2+^ uptake inhibitor (*24*) (Fig. 2F; DMSO, mean ± SD = 62.8 ± 2.1; thapsigargin, mean ± SD = 68.9 ± 3.1; p=0.1952, Kolmogorov Smirnov test). These results indicate that both active SOCE and mitochondrial Ca^2+^ entry are necessary for the observed delayed mitochondrial morphology changes. This demonstrates that ER Ca^2+^ depletion-mediated SOCE elicits a mitochondrial stress response within 10 to 30 minutes.

Cultured astrocytes express the SOCE proteins ORAI1/2/3, STIM1 and STIM2 (*4*). In alignment with the SKF-96365 experiment, we investigated whether ablating STIM1/2 proteins prevented mitochondrial fragmentation. First, we confirmed that STIM1 and STIM2 are abundantly expressed in cultured astrocytes (Fig. S2), with STIM1 being approximately 1.5 times more abundant than STIM2 (Fig. S2). Then, we isolated astrocytes from *Stim2^-/-^* mice and knocked down STIM1 protein levels via siRNA. We chose this strategy because STIM2 KO mice survive the early postnatal phase (*25*), while approximately 70% of *Stim1* KO mice die within a few hours of birth (*26*). STIM1 protein levels were reduced to approximately 20% compared to wild type levels within astrocyte cultures (Fig. S2).

Reduction of STIM1 did not lead to mitochondrial fragmentation under control conditions; however, “smaller” mitochondria were observed at baseline (Fig. 2G). Absence of STIM2 under thapsigargin did not affect mitochondrial fragmentation (DMSO, mean = 65.2 ± 1.8; TG, mean = 80.1 ± 2.3; p<0.0001, Kurskal-Wallis test with Dunn’s multiple comparisons test), while knockdown of STIM1 significantly reduced fragmentation in a similar manner to previous interventions (DMSO, mean = 59.7 ± 1.6; TG, mean=69.9 ± 2.4; p=0.054, Kurskal-Wallis test with Dunn’s multiple comparisons test) (Fig. 2G). Finally, reduction of both STIM1 and STIM2 prevented mitochondrial fragmentation under thapsigargin compared to vehicle control (DMSO, mean = 63.9 ± 1.6; TG, mean = 56.4 ± 1.5; p=0.036, Kurskal-Wallis test with Dunn’s multiple comparisons test) (Fig. 2H). In summary, changes in mitochondrial morphology were linked to the activation of SOCE, the increase in cytosolic Ca^2+^, and the entry of Ca^2+^ into mitochondria.

## Early transcriptome changes in response to ER Ca^2+^ depletion

Next, we explored transcriptome changes occurring after 60 minutes in response to either a Ca^2+^-free condition (indicating no Ca^2+^ influx) or thapsigargin-induced ER Ca^2+^ depletion, which results in long-lasting SOCE. For control, cells were kept in presence of extracellular Ca^2+^. Bulk mRNA sequencing identified approximately 17,000 different transcripts in the murine astrocytes (suppl. data 1 – TPM count table). In the Ca^2+^-free condition, compared to control cells, we observed 2,338 significantly differentially expressed genes (DEGs), which account for ∼13% of the transcriptome. Of these, 112 transcripts had a log2-fold change (log2FC) higher than 0.5, and 11 transcripts remained at a cutoff of a log2FC of 1 (Fig. 3A, B). Thapsigargin induced extensive changes in the transcriptome with 7,052 significant DEGs, in comparison to control cells, representing 41% of the transcriptome. Among these, 1,031 transcripts exhibited a log2FC higher than 0.5, and 80 exceeded log2FC of 1 (Fig. 3A, B). There were 1,606 DEGs shared between the Ca^2+^-free and thapsigargin condition (Fig. 3A). At a log2FC threshold of one, 9 transcripts overlapped between both conditions (Fig. 3C, D). Notably, five of these transcripts were cell plasticity-related transcription factors (*Csrnp1*, *Fos*, *Arc*, *Hes1*, and *Egr1*), while the remaining were signaling regulators (*Dusp1*, *Dusp6*, *Socs3*, *Gadd45g*).

**Figure 3.**
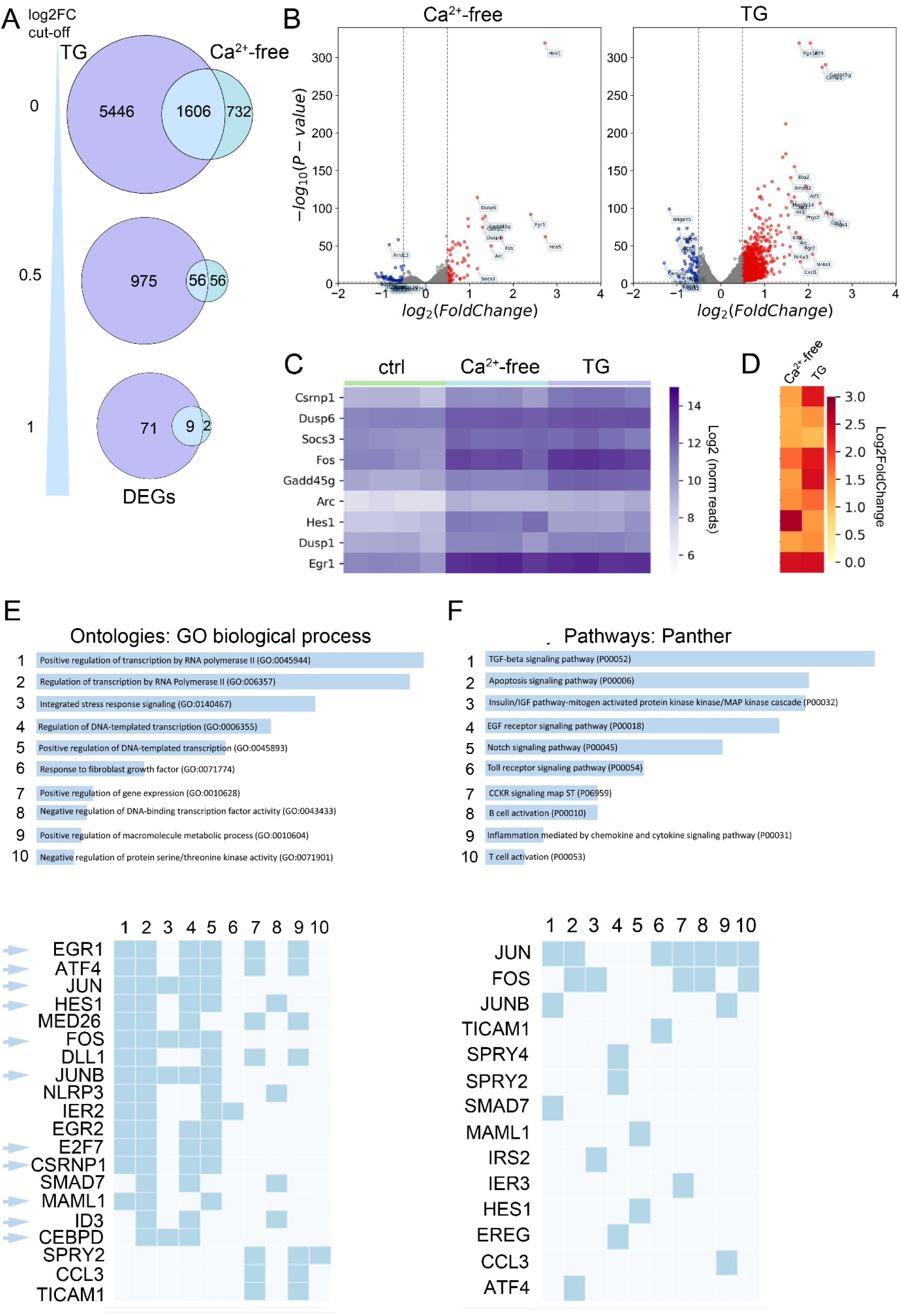
Early transcriptional changes occur in both Ca^2+^-free medium and after SERCA blockage. **A.** Venn diagram displaying gene counts and relationship of significant DEGs between cells treated with Ca^2+^-free medium and thapsigargin treated cells (TG). Data were filtered for log2FC = 0 (upper diagram), log2FC of 0.5 (middle), and log2FC = 1 (lower). **B.** Volcano plot depicting differentially expressed genes in Ca^2+^-free medium or after SERCA blockage with TG compared to untreated cells (ctrl). Significantly upregulated (red) and downregulated (blue) protein coding genes. Y-axis denotes − log10 p-adjusted values while X-axis shows log2-fold change values in gene expression. **C.** Heatmap of expression levels of the 9 most significantly regulated genes in Ca^2+^-free medium and after SERCA blockage with TG. Color indicates normalized expression reads (log2) **D.** Gene expression changes of those nine genes shown in C (log2FC values). **E - H.** Gene ontology (in D) and pathway analysis (in E). Analysis is based on 56 genes shared between both experimental conditions (Ca^2+^-free versus TG-treated, 0.5× log2FC). Numbers (1-10) link the most significant genes that are linked to corresponding biological processes or pathways. Cyan arrows point to transcription factors. Ctrl, control; DEG, differentially expressed genes; GO, gene ontology; TG, thapsigargin.

Gene ontology analysis of the 56 transcripts shared between both experimental conditions (Ca^2+^-free versus thapsigargin-treated, 0.5× log2FC) highlighted biological processes significantly associated with transcription regulation, as well as integrated stress response signaling (Fig. 3E upper panel, suppl. data 2). Pathway analysis identified signaling cascades that mostly share the same transcription factors (such as *Egr1*, *Atf4*, *Jun*, and *Hes1*) or regulators of transcription (*Med26*, *MamL1*, *ID3*) (Fig. 3F upper panel, suppl. data 3). Most of the other highly regulated transcripts were involved in cell fate differentiation (e.g., *Dll1*, *IER1*, *EGR2*, *SPRY2/4*), cell defense, innate immune signaling, or apoptosis (e.g., *Nlrp3*, *IER3*, *Ticam1*).

Under Ca^2+^-free conditions, gene ontology analysis indicated developmental processes, response to stimuli, or other top-level biological processes (Fig. 4A). This was associated with a protein interaction cluster for histone deacetylases (HDAC), which are typically involved in the activation of transcriptionally silenced chromatin, and an interaction cluster including the transcription factors *Atf4*, *Fos*, *Jun*, and *Junb* (Fig. 4C).

**Figure 4.**
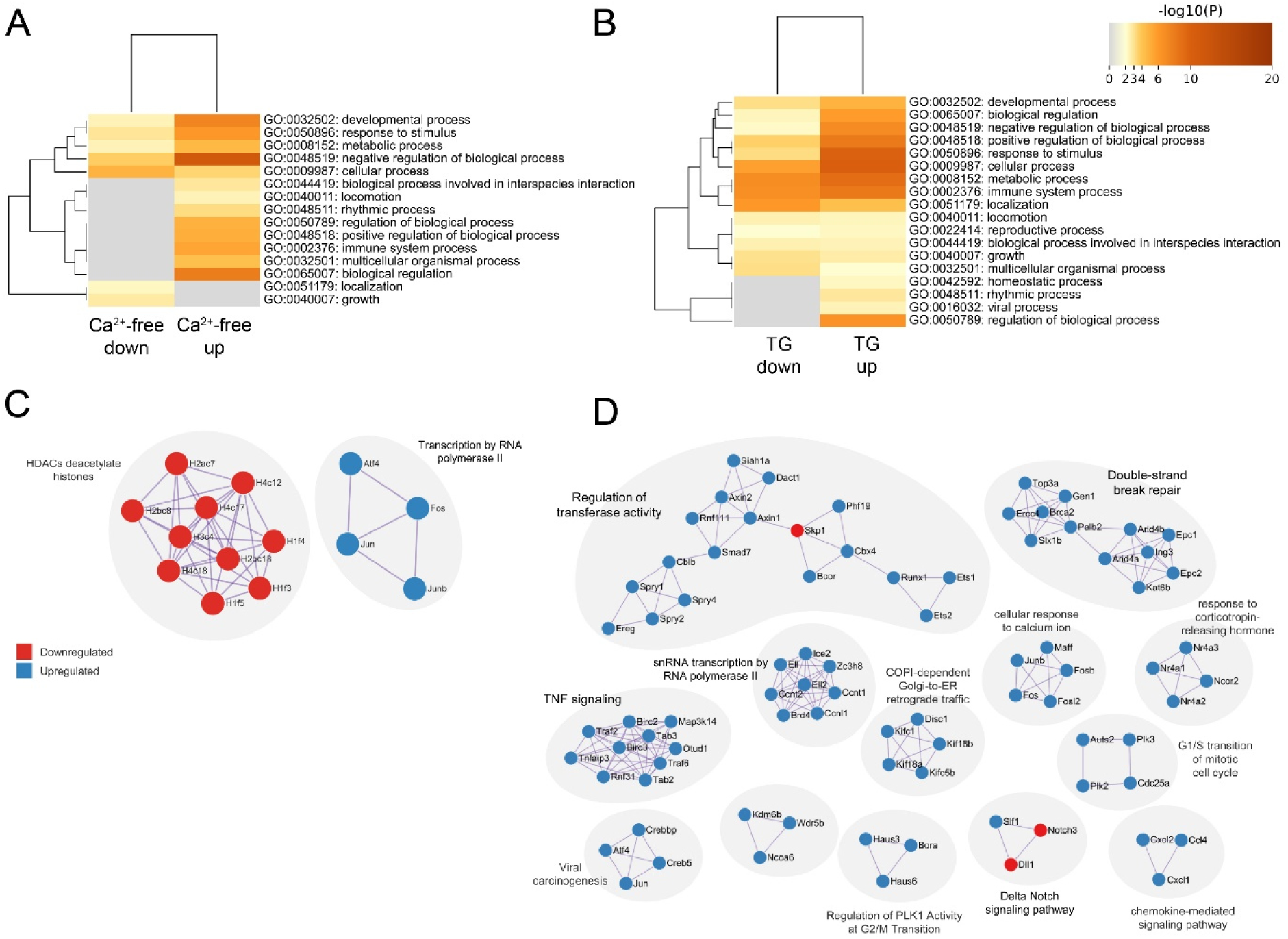
Early transcriptional changes in astrocytes after blocking Ca^2+^ entry in Ca^2+^-free medium or after SERCA blockage. Conditions: Ca^2+^-free (in A, C), thapsigargin treated (in B,D; TG) **A, B** Top level gene ontology biological processes of significantly up- and downregulated genes, colored by p-values using Metascape. **C**, **D** Gene ontology pathway analysis and protein-protein interaction computing of regulated protein coding genes. Only pathways with values of p <1×10^-6^ are shown (Bonferroni step down corrected). Top term-term interactions of regulated protein coding genes appear in corresponding clusters. Upregulated proteins are shown in blue; downregulated in red. Cluster terms (grey clouds) are based on the highest significance.

Genes specifically regulated in response to 60 minutes of thapsigargin treatment were also attributed to developmental processes or other top-level biological processes (Fig. 4B). Correspondingly, protein interaction analysis indicated the involvement in basic cellular processes like regulation of transferase activity (i.e., catalytic processes), transcription, DNA-repair, mitosis, as well as diverse signaling pathways (such as chemokine-, delta Notch-, or TNF signaling) (Fig. 4D).

## Imbalance of homeostatic SOCE leads to immediate PERK activation

In our transcriptome analysis, *Atf4* was the most prominently up-regulated factor. This immediate early transcription factor is controlled by the eukaryotic initiation factor 2 (eIF2α), which is activated by EIF2AK3, also known as PERK (*18, 27, 28*). PERK is a cell stress-related kinase that links ER Ca^2+^ activation, mitochondrial stress, and rapid transcriptome responses. Therefore, we investigated how fast PERK is activated in response to changes in Ca^2+^ homeostasis.

Astrocytes were cultured and treated with either thapsigargin or tunicamycin, a classical inducer of the ER unfolded protein response. Both thapsigargin and tunicamycin induced PERK phosphorylation at threonine 980, a commonly used marker for ER stress (Fig. 5A). PERK phosphorylation was already apparent 10 minutes after thapsigargin treatment and peaked after 30 minutes (Fig. 5B). Cells were incubated in either Ca^2+^-free medium or with the SOCE blocker SKF-96365. In both cases, PERK was phosphorylated within 10 minutes (Fig. 5C, D). Addition of SKF-96365 to thapsigargin-treated cells did not prevent PERK activation (Fig. 5C, D), although our previous experiments demonstrated that SOCE inhibition blocks mitochondrial fragmentation (see Fig. 2E).

**Figure 5.**
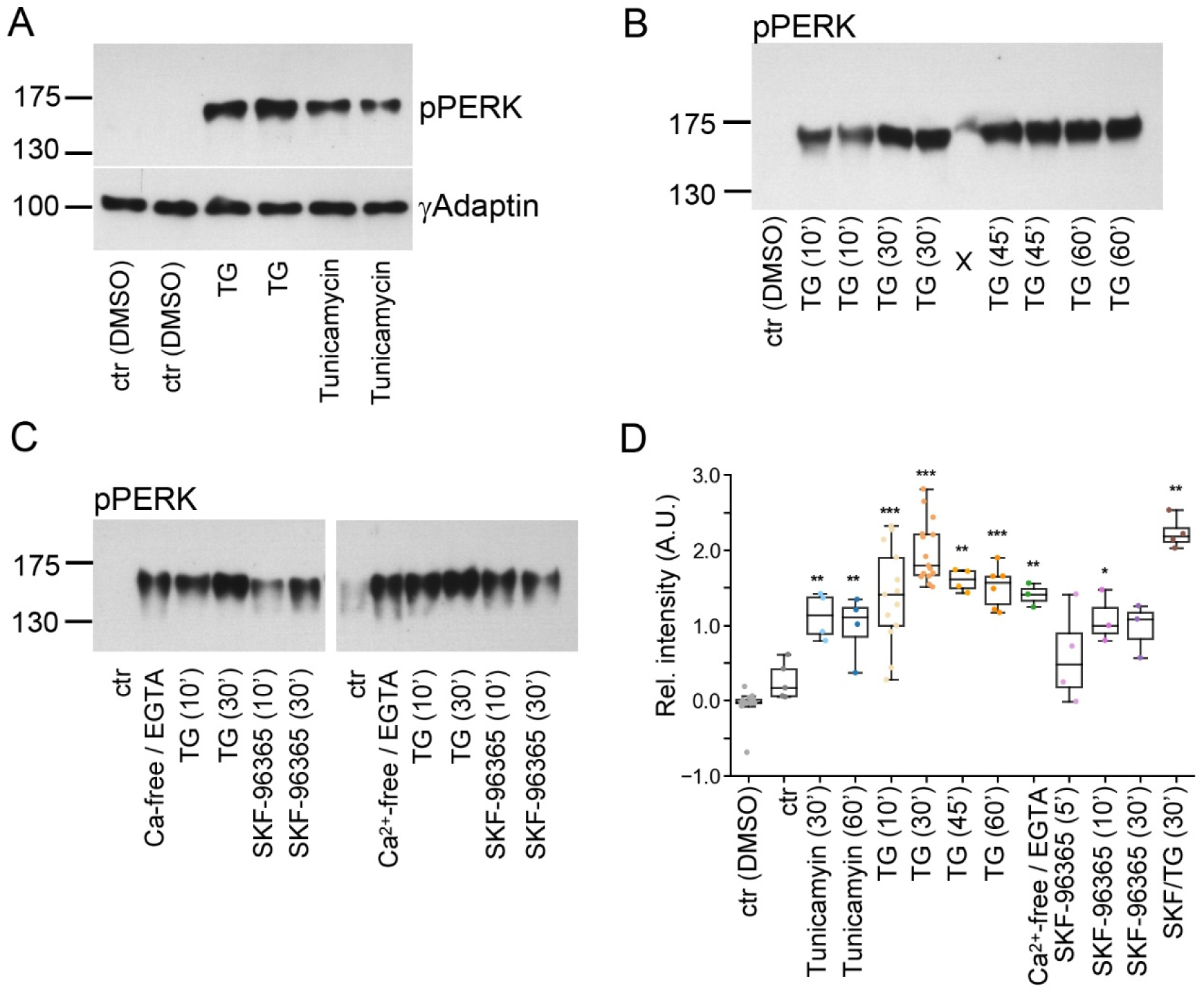
Interfering with resting SOCE causes PERK activation within minutes. Western blotting of whole cell lysates generated from astrocytes. In each lane, 40 µg protein was blotted. Due to the high molecular weight of pPERK, γ-Adaptin served as loading control (representatively shown in A). Cells were treated as indicated. **A.** PERK phosphorylation by SERCA blockage or through the unfolded protein response induced by ER stress. Cells were treated for 30 minutes with thapsigargin (TG) or tunicamycin. DMSO served as solvent control. **B.** Thapsigargin (TG) induced ER stress responses within 10 – 60 minutes. **C.** PERK phosphorylation in Ca^2+^-free medium (30 minutes), TG treatment (10 – 30 minutes), or under SOCE blockage with SKF-96365 (10 – 30 minutes). **D.** Quantification of Western blots for pPERK normalized to ctr (DMSO) with densitometry. Relative integrated densities are shown. Bar graph: mean ± SEM, overlaid with single data points. Each dot represents a biological replicate (independent cell cultures and stimulations). For each condition, data were compared to ctr (DMSO). Bonferroni adjusted p-values are indicated (*p ≤ 0.05; **p ≤ 0.01; ***p ≤ 0.001).

The E3 ubiquitin ligase Parkin is transcriptionally regulated by ATF4 in response to ER and mitochondrial stress to maintain mitochondrial integrity and protect from stress-induced cell death (*29–32*). Depending on the type, duration and severity of cellular stress, Parkin has mitophagy-dependent and mitophagy-independent stress-protective activities. Therefore, we treated astrocytes with thapsigargin or FCCP, an uncoupler of the mitochondrial proton gradient. Under vehicle control conditions (DMSO), confocal imaging confirmed tubular mitochondria and punctate endogenous Parkin immunoreactivity in astrocytes (Fig. 6A). Thapsigargin treatment caused the formation of mitochondrial spheroids and colocalization of Parkin with the mitochondrial marker Hsp60. To quantify the proximity of Parkin and Hsp60, we determined the ratio of Parkin immunoreactivity close to mitochondria (indicated in yellow in Fig. 6B). This analysis showed that under ER Ca^2+^ depletion, the localization of Parkin close to mitochondria was enhanced (Fig. 6C). A similar effect was not observed with three hours of FCCP treatment (Fig. 6C).

**Figure 6:**
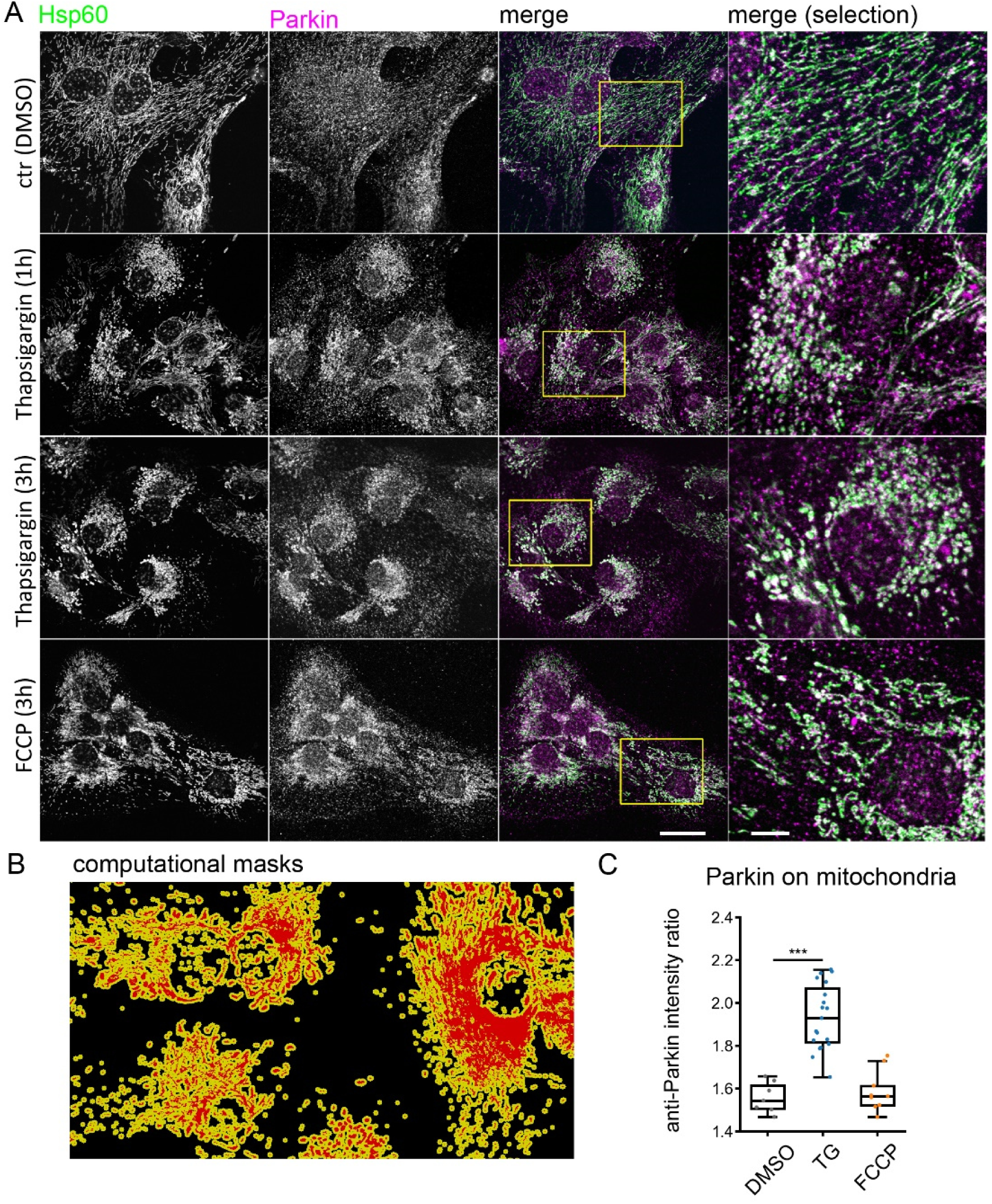
Parkin translocates to the mitochondria due to SERCA blockage-induced Ca^2+^ stress. **A** Thapsigargin (TG) treatment induces formation of mitochondrial spheroids. Immunofluorescence labeling of glial mitochondria with Hsp60 and Parkin. Experimental conditions are indicated. Confocal x.y-z images, maximum intensity projection, scale bars: 25 µm or 5 µm in detail. **B.** Computational masks were created based on Hsp60 signal. **C.** Quantification of Parkin in close proximity to mitochondria. Parkin translocation to mitochondria increases after SERCA blockage, but not in the presence of FCCP. Analysis is based on confocal images from three independent cultures (3 areas per culture). Number of cells: DMSO n = 9, TG n = 19, FCCP n = 9, t-test comparison. DMSO (vehicle) vs. TG: ***p>0.001. DMSO vs FCCP: not significant.

## Discussion

Store-operated Ca^2+^ entry (SOCE) is not only a consequence of stimulated ER Ca^2+^ depletion, but it is also present in resting cells (*3, 4, 7, 11*). We sought to determine whether homeostatic Ca^2+^ fluxes have roles beyond merely refilling cellular Ca^2+^ stores. Our data suggests that homeostatic Ca^2+^ cycling, an energy-expensive process, regulates cell health by preventing cell stress.

It is well-documented that imbalanced Ca^2+^ fluxes lead to cellular stress responses (*33, 34*). Many studies have shown that cellular stress is induced by permanent ER Ca^2+^ depletion using the SERCA blocker thapsigargin, a stimulus that unmasks the function of ER Ca^2+^ leak channels (*8, 9*). Under this condition, the depletion of the cellular Ca^2+^ store itself is thought to be the major factor for ER stress and subsequent apoptotic processes (*34*). However, thapsigargin treatment disrupts Ca^2+^ homeostasis within seconds and causes a prolonged Ca^2+^ overload in the cytosol by inducing Ca^2+^ influx from the extracellular space without refilling the ER Ca^2+^ store (*4, 10*). During sustained SOCE, blocking Ca^2+^ influx or SOCE can prevent mitochondrial fragmentation temporarily but does not prevent activation of the ER stress sensor PERK.

Primary astrocytes can manage ER Ca^2+^ loss and massive SOCE but must endure physiological consequences induced by stress sensors. The removal of SOCE under Ca^2+^-free conditions or via SERCA blockage is rapidly sensed by the cell, as indicated by PERK phosphorylation and mitochondrial fragmentation. These cellular responses are plastic responses that activate survival or death pathways. For instance, the immediate activation of PERK and the rapid upregulation of the Atf4 transcription factor are linked to processes that may rescue cells from stress. Similarly, the formation of mitochondrial spheroids, which recruits Parkin, is a biological effect typically associated with mitochondrial quality control (*35*). For instance it has been shown that Parkin can prevent thapsigargin-induced apoptosis in a mitophagy-independent manner (*30, 36*).

Removal of extracellular Ca^2+^ significantly altered the abundance of hundreds of transcripts—around 13% of the entire coding transcriptome. These acute transcriptional changes were linked to basic cellular functions such as gene expression, cell health, metabolism, and differentiation. Responses were even more pronounced under SERCA blockage, which ultimately caused cell death via the unfolded protein response (*37*). Cell stress responses after thapsigargin treatment were related but not identical to those seen when homeostatic Ca^2+^ influx is fully blocked, although both conditions are immediately sensed by PERK.

The complexity of the transcriptional responses indicates that cells expend significant energy and effort to induce a broad range of responses besides those related to cell death. For instance, the Ca^2+^-free condition causes immediate upregulation of several transcription factors involved in cell fate decisions (e.g *Hes1*, *Hes5* (*38*)) or development *(EGR2* = *Krox20*, *Dll1*, *Smad7*).

## Limitations

The first limitation of our study is that we did not perform quantitative analysis of the different Ca^2+^ responses. For the ER, we know that there is no fixed ER Ca^2+^ concentration in cultured astrocytes, with resting ER Ca^2+^ levels ranging from ‘very low’ to 150 µM of free Ca^2+^ (*4, 5*). Secondly, we anticipated seeing stronger mitochondrial Ca^2+^ responses to thapsigargin-induced SOCE; however, the responses were minimal. This might be attributed to the use of mt-pericam as a Ca^2+^-indicator, which has a Kd of 1.7 µM (*19*). We chose the mt-pericam because it can be used at a single wavelength, is well-established, and does not cause mitochondrial fragmentation or spheroid formation at high expression levels. A third limitation, as recently described for ER Ca^2+^ imaging (*4, 5*), is that the analysis of mitochondrial Ca^2+^ was dependent on ‘extreme’ low-light conditions. We believe, as discussed earlier for direct ER Ca^2+^ imaging, that this high sensitivity to light is of biological origin (*5*).

## Conclusion

We propose that the balance of Ca^2+^ fluxes between the cytosol, the ER, mitochondria, and the extracellular space, is an integrating mechanism that signals cell health. To protect cells from pathological Ca^2+^ fluxes, it may be beneficial to target late Ca^2+^ stress responses rather than immediate early Ca^2+^ fluxes in the future, as interference with either SOCE, Ca^2+^ overload, or Ca^2+^ depletion (underload), activate both beneficial and detrimental cell stress responses.

## Methods

### Agonists, antagonists, stock solutions

10 mM adenosine/water (Sigma-Aldrich, #A4036); 10 mM Btp2/DMSO; (Millipore, 203890-M); 10mM FCCP (Carbonyl cyanide-p-trifluoromethoxyphenylhydrazone)/DMSO (Sigma-Aldrich, #C2920); 25 mM SKF-96365/PBS (Abcam, #ab120280); 3 mM thapsigargin/DMSO (Abcam, #ab120286); 5mg/ml tunicamycin/DMSO (Cell signaling, #12819).

### Mouse cortical glial cells

Cortical glial cells were prepared from P3 – P5 wild type mice or STIM2 KO mice (*25*). In brief, frontal cortices from one mouse were dissected and collected in Hank’s buffered saline solution (HBSS). Then, three trituration steps were performed, and cells were cultured in basal medium (1:1 mixture of DMEM/F12 with 10% FCS, 5% horse serum, 1% penicillin/streptomycin, 0.45% glucose) containing B27 (1:50) and 10 ng/ml EGF (PeproTech). Cells were first cultured in a Poly-D L-ornithine hydrobromide (PORN) coated 75 cm^2^ culture flask at 37°C, 5% CO_2_. After 3 days, cells were washed with PBS. At day 7 in vitro (DIV7), cells were split with 20% TrpLE (Gibco) and plated at a density of 1 x 10^5^ cells/dish on poly-L-lysine-coated 10 mm glass coverslips (Marienfeld, Germany). Cells were finally used 10 – 14 DIV.

### Lentiviral vectors

For ER Ca^2+^ imaging, we used a lentiviral vector expressing ER-GCaMP6-150 (insert from RRID: Addgene #86918) under the CMV promoter (*4*). For imaging of Ca^2+^ transients in mitochondria, we cloned mt- pericam (Riken ID: RDB15144) (*19*) into the vector pSIH (CMV promoter, System biosciences). Lentiviral particles were packaged as described previously (*3*) and purified by ultracentrifugation. Glial cells were transduced 2 hours after plating on glass coverslips.

### STIM1 knockdown with siRNA

Knockdown of STIM1 mRNA was performed using FlexiTube siRNA (Qiagen) and Lipofectamine 2000 reagent (Invitrogen). Three siRNA oligos targeting different regions of the mRNA were co-transfected: Mm_STIM1_1 (5’ CAG CTT TGA GGC CGT CCG AAA ‘3); Mm_STIM1_2 (5’ CTG GTT TGC CTA TAT CCA GAA ‘3), and Mm_STIM1_6 (5’ AAG AAA GTG ATG AGT TCC TAA ‘3). As a control, the AllStars siRNA control nucleotide (Qiagen) was used. For transfection, 4 µl Lipofectamine 2000 and 20 pM siRNA were incubated for 30 minutes in 500 µl serum-free OptiMEM (Gibco). The mixture was then added to 5x 10^5^ glial cells in 1.5 ml medium (antibiotic-free). Further cultivation medium was added after 4 to 6 hours, and the whole medium was replaced the next day. Subsequent experiments were performed at 48 to 72 hours after transfection.

### Ca^2+^ imaging: ER, cytosol

*Imaging solutions:* HEPES-buffered ACSF (artificial cerebrospinal fluid, 120 mM NaCl, 2.5 mM KCl, 1.2 mM MgCl_2_, 2.4 mM CaCl_2_, 1.2 mM NaH_2_PO_4_, 26 mM NaHCO_3_, 10 mM Glucose, 10 mM HEPES); Ca^2+^-free HEPES-buffered ACSF (123.6 mM NaCl, 2.5 mM KCl, 1.2 mM MgCl_2_, 1.2 mM NaH_2_PO_4_, 26 mM NaHCO_3_, 10 mM Glucose, 10 mM HEPES, 0.1 mM EGTA).

During imaging, cells were continuously perfused with a Minipuls 3 peristaltic pump (speed of the perfusion pump: 10 A.U.; purple tubing, Gilson). Solution was pre-warmed (37°C) with an in-line heater system (Warner) and a stage heater (37°C) (Luigs & Neumann). Agonists and antagonists were applied with the perfusion system. Extreme low light conditions were necessary (see (*4, 5*)).

Dual color Ca^2+^ imaging (Fig. 1B, ER-cytosol) was performed using the genetic ER Ca^2+^ indicator ER- GCaMP6-150 and the red fluorescent synthetic Ca^2+^ indicator Cal-590 acetoxymethyl ester (AAT Bioquest) as previously described (*4*). Cytosolic calcium imaging (Fig. S1) was performed with Oregon Green BAPTA1-AM as described (*4*).

### Imaging of mitochondria (mt-pericam and Mitotracker CMXros)

CMXros (MitoTracker Red, Invitrogen, M7512)) was diluted to a 5 mM stock solution with DMSO and stored at -20°C. For mitochondria imaging under live cell conditions, cells were stained with 25 nM dye in medium at 37°C for 10 minutes. Imaging was started after one wash in HEPES-ACSF. For epifluorescent imaging, an upright fixed microscope (BXWI, Olympus) equipped with a CoolLED (Visitron Systems) and a X-cite 120Q excitation light source (Lumen dynamics) was used in combination with an UPLSAPO 60xW/N.A. 1.0 objective. Images were captured using the Rolera-XR camera (Qimaging) operated by the Stream Pix 4.0 software (Norpix) with the settings: 696 x 520 pixels, 2 Hz, gain 17.800, offset 0. CMXros was excited with 550 nm wavelength (CoolLED, emission filter: band pass 545/30 nm / 570 nm / 620/60 nm; DsRed filter set, AHF). Thapsigargin (3 µM) or vehicle were applied by continuous perfusion in HEPES-ACSF at 37°C. For analysis of mitochondrial morphology, up to 20 randomly chosen areas with widespread cell coverage were imaged within 10 minutes. For quantification of mitochondrial fragmentation, images were analyzed by NIH ImageJ software (Rasband, W.S., ImageJ, U.S. National Institutes of Health, Bethesda, Maryland, USA, https://imagej.nih.gov/ij/) (68). As shown in supplemental figure 2, 8-bit raw images were inverted, sharpened, and converted to a binary code by adjusting the threshold (Dagda et al., 2009). Four different size categories were chosen for particle analysis via the particle analyzer plugin in ImageJ, namely: 5 – 15 pixel2, 16 – 50 pixel2, 51 – 100 pixel^2^, and 101 to infinity pixel^2^. To illustrate this categorization in the mitochondrial network, a colored mask image was produced showing the particle sizes in different color codes. Data were plotted showing the mean particle size per category ± SD for each incubation time point (Fig. 2B) and as area corrected values for the category 5-15 pixel^2^ (Fig. 2C- H).

For qualitative imaging of mitochondrial Ca^2+^ (Fig. 1D), ratiometric pericam targeted to the mitochondrial matrix (mt-pericam) (*19*) was used as a single-wavelength indicator and was excited at 380 nm. Ratiometric pericam has a bimodal excitation spectrum peaking at 415 and 494 nm, whereas signal changes are more pronounced when pericam is excited at 380 – 410 nm (*19*). The GECI shows monophasic Ca^2+^ dependence (Kd ∼ 1.7 µM). Emission intensities in response to 380 nm excitation are reduced when Ca^2+^ binds to mt-pericam. This strategy allowed imaging of qualitative changes of mitochondrial Ca^2+^, with 5 Hz, under continuous monitoring in low light conditions.

### Indirect immunofluorescence and confocal microscopy

Glial cells were fixed with PBS-buffered 4% paraformaldehyde (pH 7.4) for 15 minutes at 37°C. Blocking solution was PBS with 10% goat serum, 0.1% Triton X100, and 0.1% Tween 20. Antibodies were diluted in blocking solution. Coverslips were washed with 0.1% Triton X100, 0.1% Tween 20 in PBS. Primary antibodies were incubated in blocking solution for 1 to 3 hours. Fluorochrome-conjugated secondary antibodies were used for 1 hour. Cell nuclei were labeled with DAPI (2 mg/ml stock solution, freshly diluted 1:5000 in PBS) for 5 minutes. Following DAPI labeling, cells were washed twice with PBS and coverslips were mounted with Aquapolymount (Polysciences).

*Primary antibodies:* goat anti-HSP60 (N-20) (1:100), Santa Cruz (#Sc-1052; RRID: AB_631683); mouse anti-Parkin (PRK8) (1/200), Santa Cruz (#Sc-32282; RRID: AB_628104).

*Secondary antibodies:* donkey anti-goat Alexa 488 (1:800) Invitrogen (A-11055; RRID: AB_2534102); donkey anti-mouse Cy3 (0.5mg/ml) (1:800), Jackson (#715-165-151; AB_2315777).

Images were acquired using an inverted IX81 microscope equipped with an Olympus FV1000 confocal laser scanning system, a FVD10 SPD spectral detector and diode lasers of 405, 473, 559 and 635 nm. All images shown were acquired with an Olympus UAPO 20x (air, numerical aperture 0.70) or UPLSAPO 60x (oil, numerical aperture:1.35) objective. For high-resolution confocal scanning, a pinhole setting representing one Airy disc was used. In case of high-resolution imaging, confocal settings were chosen to meet an optimum resolution of at least 3 pixels per feature in the x-y direction. In the z-direction, 300 nm steps were used. 12-bit z-stack images were processed by maximum intensity projection and were adjusted in brightness and contrast using ImageJ software (Rasband, W.S., ImageJ, U.S. National Institutes of Health, Bethesda, Maryland, USA, https://imagej.nih.gov/ij/) (68). Images are shown as RGB images (8-bit per color channel). Fluorescence images were processed for final presentation using Adobe Photoshop CS5.

### RNA sequencing

Glial cells were incubated for 60 minutes in medium with 2mM CaCl_2_ (control), Ca^2+^-free medium, or in medium with 2mM CaCl_2_ and 3 µM thapsigargin. Cells were lysed lysed in RLT buffer with 1% β-mercaptoethanol, and homogenized with a 25G syringe. RNA quality was checked using a 2100 Bioanalyzer with the RNA 6000 Nano kit (Agilent Technologies). The RIN for all samples was >9.3. DNA libraries suitable for sequencing were prepared from 500 ng of total RNA with oligo-dT capture beads for poly-A-mRNA enrichment using the TruSeq Stranded mRNA Library Preparation Kit (Illumina) according to the manufacturer’s instructions. Sequencing of pooled libraries, spiked with 1% PhiX control library, was performed at ∼30 million reads per sample in single-end mode with 75 nt read length on the NextSeq 500 platform (Illumina) with 1 High Output Kit v2.5. Demultiplexed FASTQ files were generated with bcl2fastq2 v2.20.0.422 (Illumina). Illumina reads were quality- and adapter-trimmed via Cutadapt (*39*) version 2.5 using a cutoff Phred score of 20 in NextSeq mode. Reads without any remaining bases were discarded (command line parameters: --nextseq-trim=20 -m 1 -a AGATCGGAAGAGCACACGTCTGAACTCCAGTCAC). Processed reads were mapped to the mouse genome (GRCm38.p6 primary assembly and mitochondrion) using STAR v2.7.2b with default parameters based on RefSeq annotation version 108.20200622 for GRCm38.p6 (*40*). Read counts on the exon level summarized for each gene were generated using featureCounts v1.6.4 from the Subread package (*41*). Multi-mapping and multi-overlapping reads were counted strand-specific, and reversely stranded with a fractional count for each alignment and overlapping feature (command line parameters: -s 2 -t exon -M -O --fraction). Raw read counts per gene were converted to Transcripts Per Million (TPM) values based on combined exon length.

### RNA-Seq data analysis

The count output was utilized to identify differentially expressed genes using DESeq2 (*42*) version 1.24.0. Read counts were normalized by DESeq2 and fold-change shrinkage was applied by setting the parameter “betaPrior=TRUE”. Differential gene expression was analyzed for genes with an adjusted p-value (padj) after Benjamini-Hochberg correction < 0.05 and at a log2-fold change of ≥ 0.5. Only protein-coding genes were considered for analysis of differentially expressed genes (DEG). Venn diagrams were created with SRplot (SRplot- Science and Research online plot) (*43*). Volcano plots and heatmaps were created using Python version 3.12 (bioinfokit.visuz and seaborn). Ontologies and pathways were analyzed and visualized as bar graphs and clustergrams with enrichr (https://maayanlab.cloud/Enrichr/),(44). Enrichment heatmaps and protein-protein interaction networks (PPI) were generated with Metascape (http://metascape.org) (*45*) and PPIs were adapted in Cytoscape (*46*).

### Western blot analysis

Glial cells (on 30 mm dish, poly-L-lysin coated) were lysed at DIV 6 – 7 using a cell scraper and 200 µl cold lysis buffer (1% NP40, 50 mM HEPES pH 7.5, 150 mM NaCl, 10% Glycerol, 1 mM sodium fluoride, 10 mM sodium pyrophosphate, 2 mM sodium orthovanadate, 5 mM EDTA, supplemented with one EDTA-free protease inhibitor mini tablet / 5 ml of buffer (Roche Cat#4693159001). Lysates were incubated on ice for 15 minutes. Cells were sonicated twice (Hielscher sonifier UP50, M1 sonotrode, 80% power, 10 cycles at 0.5 s) and placed back on ice for 10 minutes. Lysates were centrifuged for 5 minutes at 4°C at 15,000xg. Protein concentration of the samples was determined by Pierce BCA protein assay kit (Thermo Scientific). After standard SDS-PAGE, 40 µg of total protein per lane was transferred onto PVDF membranes (Immun-Blot, BioRad) using a semi-dry blotter (BioRAD). After blocking for 30 minutes at RT, primary antibodies were incubated for 3 hours at RT. Blots were developed on X-ray films (Fujifilm Super RX) using ECL (Immobilon Western HRP Substrate, Merck Millipore). For quantification, X-ray films were first photographed with a 12MP Canon camera on a white transilluminator plate (PeqLab Gel Documentation system). Protein bands were quantified using ImageJ (Rasband, W.S., ImageJ, U.S. National Institutes of Health, Bethesda, Maryland, USA, https://imagej.nih.gov/ij/) (*47*).

*Primary antibodies:* rabbit anti-phospho-PERK (Thr980) (16F8) (1:2,000) Cell signaling (#3179; RRID: AB_2095853): mouse anti-γ-Adaptin (clone 88) (1:2,000) BD biosciences (#610385; RRID: AB_397768).

S*econdary antibodies:* Peroxidase AffiniPure Goat Anti-Mouse IgG (H+L) (1:5,000), Jackson, (# 115-035- 146; RRID: AB_2307392); Peroxidase AffiniPure Goat Anti-Rabbit IgG (H+L) (1:5,000), Jackson (# 111- 035-144; RRID: AB_2307391).

### Statistics

Data were tested for significance using Python 3 with packages scipy.stats (*48*) and GraphPad Prism software v10.4.0. Normal distribution was assessed with the Shapiro test. When outside of normal distribution, data were tested for significant differences with the two-sided Mann-Whitney U test. Normally distributed data were initially tested for equal variance with the Bartlett’s test. Two groups with unequal variance were compared with the Welch’s t-test, while those with equal variance with an unpaired, two-tailed t-test. Data are presented as the mean ± standard error of the mean (s.e.m.). A p-value adjusted < 0.05 was considered statistically significant. In boxplots, single data points are shown. The boxes extend from the 25th to the 75th percentile and lines within boxes show the median. The whiskers extend from the smallest to the highest value except for outliers. Other graph characteristics can be found in the figure legends.

## Data availability

The RNA-seq data are available at NCBI GEO: GSE242678. All other data needed to evaluate the conclusions in the paper are present in the paper or the Supplementary Materials.

## Acknowledgements

We thank Margarete Göbel of the Core Unit SysMed at the University of Würzburg for excellent technical support. We are grateful for excellent technical assistance by Michaela Kessler. We would like to thank Takeharu Nagai and Atsushi Miyawaki for sharing mt-pericam, and Timothy A. Ryan (Weill Cornell Medical College, New York) for sharing ER-GCaMP6–150.

## Author contributions

Conceptualization: CF, AS, KW, RB; Acquisition of data: CF, AS, F. Schmitt, RB; Methodology: JJ, FS, IL; Investigation and formal analysis: CF, AS, F. Schlott, TB, RB; Resources: BN; Supervision: RB; Visualization: CF, AS, F. Schlott, RB; Writing – Original Draft: CF, RB; Writing – Review and Editing: all authors; Funding acquisition: AS, RB.

---

**Figure S1.**
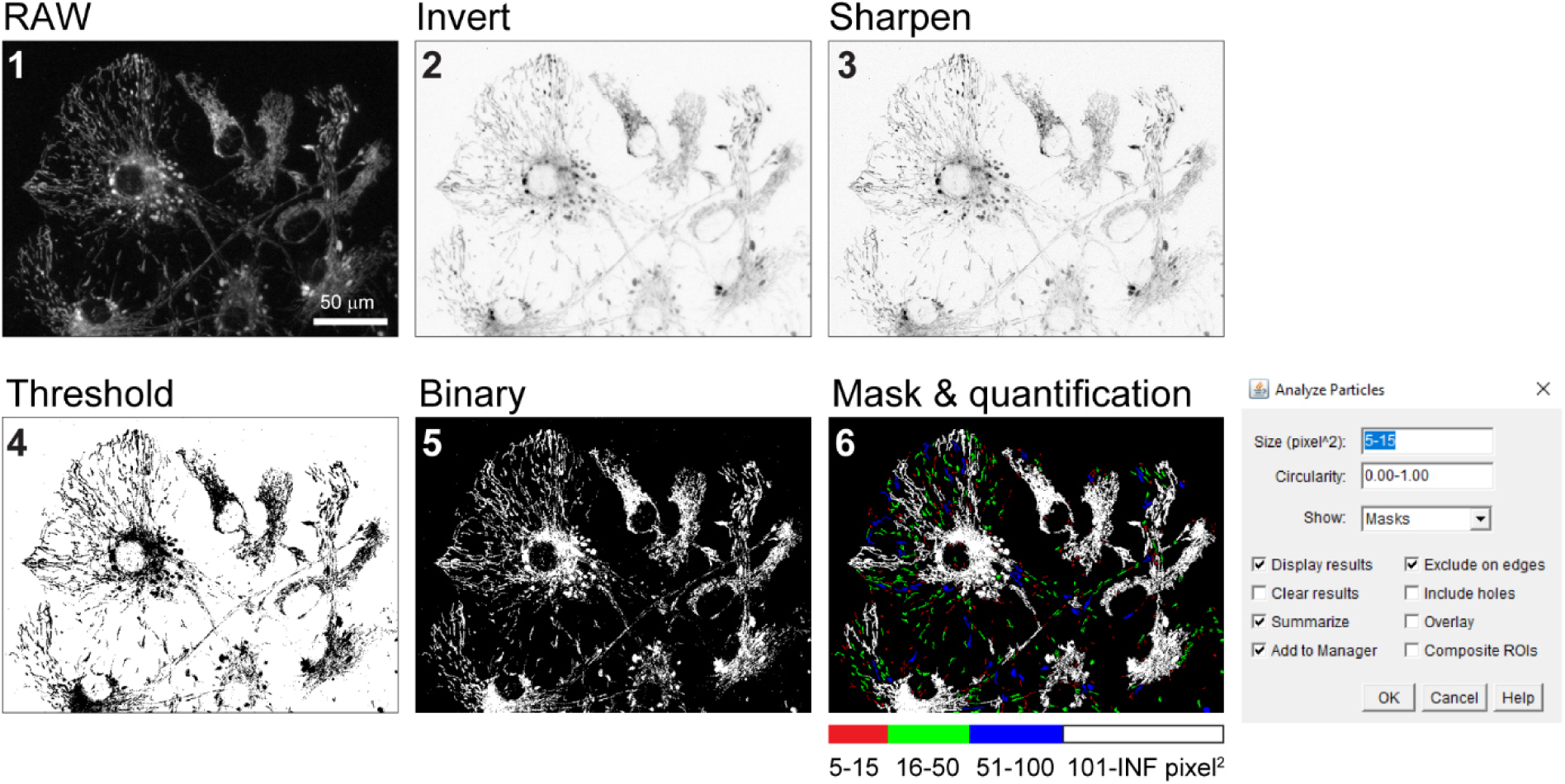
Quantification of mitochondrial fragmentation. Image raw data (8-bit, 1) were inverted and sharpened (2, 3). Signal thresholds (4) were determined to compute binary masks (5) for quantification of mitochondrial particles by size category (6).

**Figure S2.**
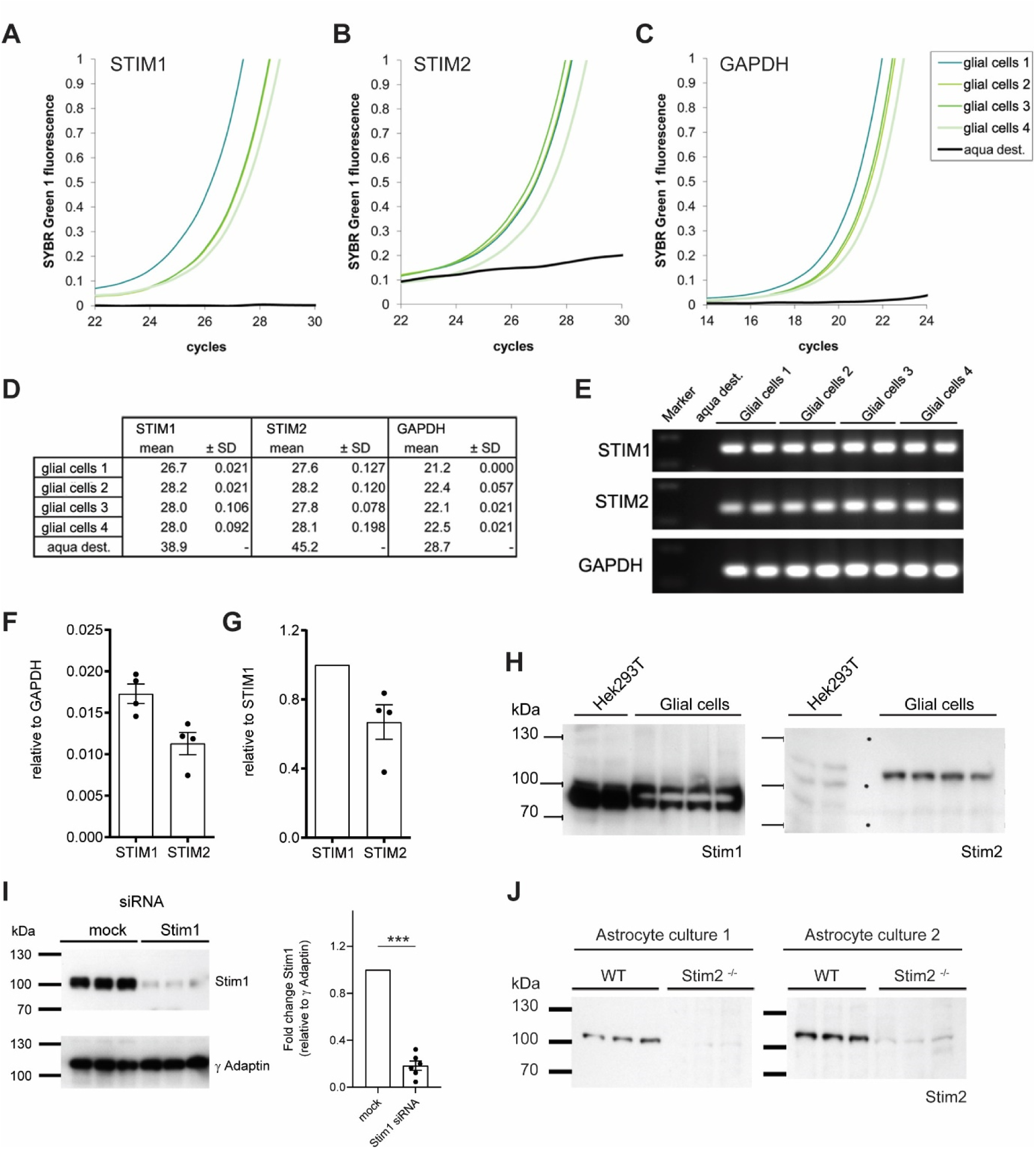
STIM1 and STIM2 abundance in glial cells, and confirmation of their knock-down and knock-out in culture. **A-C.** SYBR green fluorescence traces from qPCR amplification of STIM1 (A), STIM2 (B), and GAPDH (C) in four glial RNA samples from independent cultures. **D**. Quantification of CT values, **E**. visualization of PCR products, and **F, G**. relative quantification of mRNA abundance relative to GAPDH (F) and STIM1 (G). n= 4 biological replicates, Mean ± SEM. **H**. Western blots probing Stim1 and Stim2 proteins in Hek239T cells and primary glial cultures (n= 4 biological replicates). **I**. Western blot and quantification of siRNA mediated knockdown of Stim1 in glial cultures (n= 2 biological replicates measured as technical duplicates). γ Adaptin was used as loading control. Unpaired, two-tailed t-test: t(10) = 20.7, p<0.001, ***. **J**. Western blots from two independent glial cultures generated from Stim2 ^-/-^ pups and Stim2 ^+/+^(WT) littermates.

